# Cloudberry-derived nanovesicles: *in vitro* functional effects in skin cell models and characterization of molecular cargo

**DOI:** 10.64898/2026.07.28.741293

**Authors:** Ramila Mammadova, Feby Wijaya Pratiwi, Carmen Laezza, Keerthanaa Balasubramanian Shanthi, Dávid Papp, Carmina Sirignano, Dilki Madubhashani, Maria Manuela Rigano, Gitta Schlosser, Seppo Vainio

## Abstract

Cloudberry (*Rubus chamaemorus* L.)-derived nanovesicles (NVs) represent a promising but still poorly characterized class of plant-derived vesicles with potential relevance for skin-related applications. Here, we isolated cloudberry fruit-derived NVs and investigated their physicochemical and molecular properties, cellular uptake, cytocompatibility, and functional effects in human dermal fibroblasts (HDF) and HaCaT keratinocytes. Nanoparticle tracking analysis and transmission electron microscopy confirmed a nanosized vesicle preparation with characteristic round morphology, while protein quantification supported reproducible isolation of NV-associated material. In vitro, cloudberry NVs showed concentration-dependent effects on cell viability and proliferation, with lower doses being better tolerated. Labelled NVs were internalized by both HDF and HaCaT cells in a time-dependent manner. Under oxidative stress conditions, cloudberry NVs reduced H_2_O_2_-induced senescence-associated β-galactosidase staining in HDFs and exerted cytoprotective effects in both cell lines, alongside measurable cell-free antioxidant activity in the DPPH assay. In scratch wound-healing assays, cloudberry NVs modulated wound closure in a dose-dependent manner, with the lowest tested concentration showing the most favorable response. UHPLC-MS/MS-based proteomics and metabolomics further indicated the presence of diverse secondary metabolites and stress-related protein cargo. Together, these results support the view that cloudberry-derived NVs are biologically active plant nanovesicles with potential utility in skin-related regenerative applications.

**Graphical Abstract:** 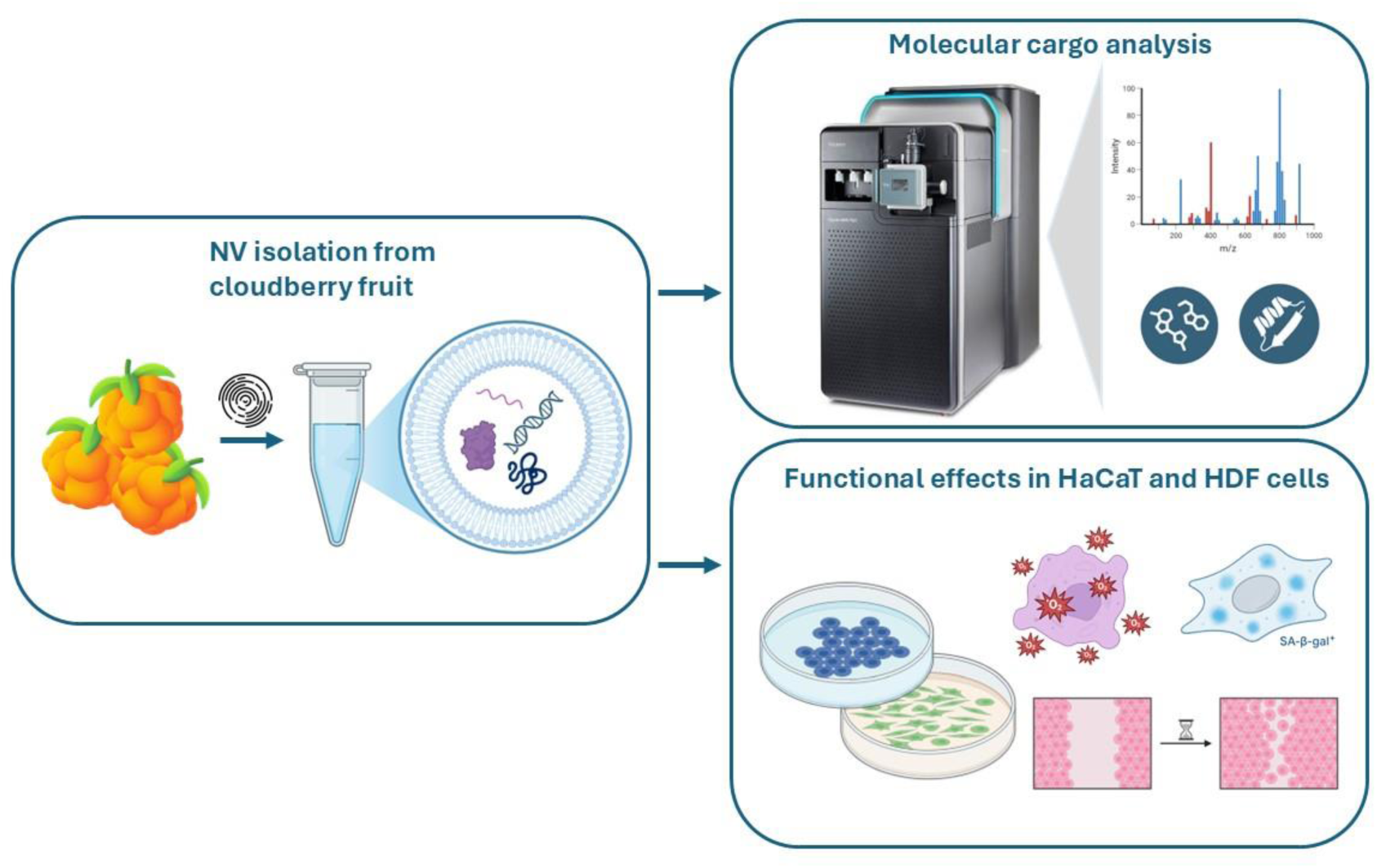

## 1. Introduction

Extracellular vesicles (EVs) are membrane-enclosed nanoscale particles naturally released by cells and are widely recognized as important mediators of intercellular communication through the transfer of proteins, lipids, nucleic acids, and metabolites ^1^. Recent MISEV2023 recommendations have further emphasized the importance of careful terminology, isolation, and characterization when describing EV-related particles and vesicle preparations ^1,2^. In recent years, increasing attention has also been given to plant-derived extracellular vesicles, or plant-derived nanovesicles (PDNVs), as a promising class of naturally derived vesicular structures with potential biomedical relevance ^3,4^.

Plant-derived extracellular vesicles have attracted growing interest because they combine natural origin with biological activity, relative stability, and favorable biocompatibility ^3,4^. Recent studies and reviews highlight these vesicles as promising candidates for therapeutic and regenerative applications because they carry diverse bioactive cargo and can modulate key cellular processes such as proliferation, migration, inflammation, and oxidative stress responses, with potential relevance for wound healing and skin regeneration ^5–8^. For example, wheat-derived nanovesicles were shown to promote proliferation and migration in wound-healing models ^6^, ginger-derived nanoparticles demonstrated anti-inflammatory activity in intestinal models ^5^, and plant-derived extracellular vesicle preparations have shown antioxidant and reparative effects in human skin fibroblasts ^7^.

The skin is continuously exposed to environmental and endogenous stressors, and oxidative stress is a major contributor to skin aging, cellular dysfunction, and impaired tissue repair ^9^. Because of this, there is increasing interest in identifying natural vesicle-based systems that could protect skin cells, reduce stress-associated damage, and support wound-healing-related responses ^9^.

Berry-derived materials are especially attractive in this context because berries are rich in polyphenols, vitamins, lipids, and other phytochemicals associated with antioxidant and health-promoting properties ^10^. Cloudberry (*Rubus chamaemorus* L.), an Arctic berry valued both nutritionally and traditionally, is recognized as a rich source of beneficial antioxidant bioactive compounds ^10^. However, while the fruit itself has been studied for its nutritional and bioactive composition, much less is known about nanovesicles derived from cloudberry and their possible biological effects in skin-related models. Despite the rapid expansion of the plant vesicle field, important knowledge gaps remain regarding vesicles isolated from specific plant sources, including their physicochemical characteristics, molecular cargo, cellular uptake, and biological effects in relevant in vitro systems ^3,4,9,11^. Notably, our recent work demonstrated that cloudberry-derived nanovesicles remain stable under simulated gastrointestinal conditions, are efficiently taken up and transported across Caco-2 cell monolayers without cytotoxicity, and show favorable biodistribution and low immunogenicity in vivo, further supporting their promise as biologically relevant plant-derived nanosystems ^12^. In the case of cloudberry-derived nanovesicles, information is still very limited, particularly regarding their interaction with skin-related cells and their potential effects on oxidative stress, senescence, and wound-healing-associated processes.

Therefore, the aim of the present study was to isolate and characterize cloudberry fruit-derived nanovesicles and to investigate their biological effects in skin-related in vitro models. Specifically, we evaluated their cytocompatibility, cellular uptake, antioxidant and anti-senescent potential, and wound-healing-related activity, and further explored their molecular composition by metabolomic and proteomic analyses.

## 2. Materials and Methods

### 2.1. Isolation of cloudberry-derived nanovesicles

Cloudberry (*Rubus chamaemorus L.*) was purchased from a local store in Oulu, Finland, and NVs were isolated using dUC as previously described ^12^.

### 2.2. Physicochemical characterization

#### 2.2.1. Nanoparticle Tracking Analysis (NTA)

The particle size distribution and concentration of cloudberry-derived nanovesicles were determined by nanoparticle tracking analysis (NTA) using a NanoSight NS300 instrument (Malvern Panalytical, Malvern, UK). For each measurement, 1 µg of NVs, calculated based on protein content, were diluted to a final volume of 1 mL with Milli-Q water, and the entire diluted sample was loaded into the instrument. Measurements were performed using a camera level of 14 and a detection threshold of 3. The resulting data were then analyzed with NTA software version 3.4.

#### 2.2.2. Qubit Protein Assay

The protein concentration of NVs was determined using the Qubit Protein Assay Kit and measured with a Qubit 3.0 Fluorometer (Thermo Fisher Scientific, Rockford, IL USA).

#### 2.2.3. Transmission Electron Microscopy (TEM)

The surface morphology of cloudberry NVs was examined by transmission electron microscopy (TEM) using a Tecnai G2 Spirit instrument (FEI, Eindhoven, The Netherlands). Briefly, 2 µL of each sample was deposited onto a glow-discharged Formvar carbon-coated grid. The samples were then negatively stained with 2% uranyl acetate and imaged using the Tecnai G2 Spirit transmission electron microscope. Micrographs were acquired at a magnification of 1:23,000 with a charge-coupled device camera (Quemesa; Olympus Soft Imaging Solutions GmbH, Münster, Germany).

### 2.3. In vitro characterization

#### 2.3.1. Cell cultures

Human dermal fibroblasts, adult (HDF-Ad; CLS Cell Lines Service GmbH, Cat. No. 300606), were cultured in Minimum Essential Medium (MEM) with Earle’s salts and stable glutamine (Biowest, Cat. No. L0416-500), supplemented with 10% fetal bovine serum (FBS) (Gibco) and 1% penicillin–streptomycin (Sigma), at 37°C in an incubator containing 5% CO₂.

HaCaT keratinocytes were cultured in DMEM/F-12 (1:1) with GlutaMAX (Gibco, Cat. No. 31331-028), supplemented with 10% fetal bovine serum (FBS), at 37°C in an incubator with 5% CO₂.

#### 2.3.2. Cytotoxicity and cell proliferation assays

An IncuCyte cell proliferation assay and MTT assay were used to evaluate the cytotoxicity, cell viability, and proliferation of cells treated with cloudberry-derived nanovesicles (NVs).

##### 2.3.2.1. IncuCyte cell proliferation assay

For the IncuCyte cell proliferation assay, cells were detached with trypsin, collected by centrifugation at 300 × g for 3 min, counted using a TC20™ Automatic Cell Counter (Bio-Rad), and resuspended in culture medium supplemented with 10% FBS. Cells were seeded in 100 µL medium into 96-well flat-bottom plates at densities of 2 × 10^3^ cells/well for HDFs and 5 × 10^3^ cells/well for HaCaT cells, and incubated overnight at 37 °C in 5% CO_2_. After overnight attachment, the medium was removed, the cells were washed with phosphate-buffered saline (PBS), and fresh medium containing cloudberry NVs was added. SYTOX Green high-affinity nucleic acid stain, diluted 1:10,000 in culture medium, was used to label dead or dying cells. Plates were scanned every 3 h using the IncuCyte live-cell imaging system at 10× magnification with the standard scan setting. Dead or dying cells were automatically quantified using the IncuCyte image analysis software. All samples were analyzed in triplicate.

##### 2.3.2.2. MTT assay

Cell viability was additionally assessed using the MTT assay. Cells were seeded into 96-well plates at the same densities used for the IncuCyte assay (2 × 10^3^ HDFs/well and 5 × 10^3^ HaCaT cells/well) and allowed to attach overnight. After 48 h of treatment, the medium was removed and 50 µL of diluted MTT solution was added to each well. After 3 h incubation, the solution was removed carefully and the resulting formazan crystals were dissolved in 100 µL of isopropanol solution. Absorbance was recorded at 570 nm using a microplate reader.

#### 2.3.3. Cellular uptake of cloudberry NVs by the cells

HaCaT keratinocytes were seeded onto glass coverslips placed in 12-well plates at different densities depending on the incubation period: 1 × 10^5^ cells/well for the 24 h time point, and 2 × 10^5^ cells/well for the 2, and 8 h time points. Human dermal fibroblasts were seeded on coverslips in 24-well plates at densities adjusted according to the experimental time points: 2 × 10^4^ cells/well for 24 h incubation and 5 × 10^4^ cells/well for 2 and 8 h incubations. Cloudberry-derived NVs were labelled with green lipophilic tracer DiOC_18_(3) (3,3’-Dioctadecyloxacarbocyanine perchlorate) (Invitrogen, USA). After attachment, the cells were treated with labelled NVs at a final concentration of 10 µg/mL in complete culture medium. Control cells were treated with medium alone. Cells were incubated with labelled CB NVs at 37 °C for 2, 8, and 24 h. After incubation, the cells were washed with PBS, stained with LysoTracker Red DND-66 (Invitrogen, USA) for lysosomal staining, and fixed with 4% paraformaldehyde (PFA). The fixed cells were then stained with a blue fluorescent dye Hoechst 33258 (Invitrogen, USA) for nuclear visualization. The coverslips were mounted on the glass slide and the results were visualized using a Leica Stellaris 8 DIVE confocal and multiphoton microscope (Leica Microsystems GmbH, Germany).

#### 2.3.4. Senescence-associated β-galactosidase staining assay

Anti-senescent activity was evaluated using the Cell Senescence β-Galactosidase Staining Kit (HY-K1089, MedChemExpress, Monmouth Junction, NJ, USA) according to the manufacturer’s instructions. Briefly, HDF cells were seeded onto coverslips in 24-well plates at a density of 1×10^5^ cells/well and allowed to attach overnight. The cells were washed with PBS and pretreated with cloudberry NVs at 2, 5, or 10 µg based on protein content for 24 h before induction of senescence with 250 µM H_2_O_2_. After exposure to H_2_O_2_ for 1.5 h at 37°C, the cells were washed with PBS and incubated with fresh medium for 20 min at 37°C. The cells were then fixed with β-galactosidase staining fixative solution at room temperature, washed three times with PBS, and incubated overnight at 37°C in a CO_2_-free incubator with freshly prepared staining working solution. After staining, the cells were washed with PBS and examined under a Zeiss Axio Imager 2 motorized microscope equipped with an Axiocam 506 color camera (Carl Zeiss Microscopy GmbH, Jena, Germany). SA-β-Gal-positive cells and total cells were calculated from the brightfield images using FiJi/ImageJ software (version 1.54f), and the percentage of SA-β-Gal-positive cells was calculated for each image.

#### 2.3.5. Scratch wound-healing assay

A scratch wound migration assay was performed to evaluate the effect of cloudberry NVs on cell migration in vitro. HDF and HaCaT cells were seeded in Incucyte ImageLock 96-well plates at densities of 2 × 10^4^ and 3 × 10^4^ cells/well, respectively, and cultured until a confluent monolayer was formed. Once full confluence was reached, the medium was removed and uniform wounds were generated using the Incucyte WoundMaker device (Essen Bioscience, San Jose, CA, USA). After scratching, the wells were washed with PBS to remove detached cells, and serum-free medium containing cloudberry NVs at 2, 5, or 10 µg based on protein content was added. Wound closure was monitored for 48 h using the Incucyte S3 live-cell analysis system (Essen Bioscience), and digital images were acquired every 2 h. The remaining wound confluence (%) was quantified over time to assess cell migration. Assays were performed in triplicate.

#### 2.3.6. Antioxidant activity

##### 2.3.6.1. 1,1-Diphenyl-2-picrylhydrazyl (DPPH) Assay

The assay was performed with modifications based on a previously published method^13^. Briefly, cloudberry NVs at 2, 5, and 10 µg, based on protein content, were adjusted to a final volume of 10 µL with buffer and mixed with 90 µL of 0.2 mM DPPH solution. A control sample was prepared by mixing 10 µL of Tris-HCl buffer with 90 µL of DPPH solution. The reaction mixtures were incubated at room temperature in the dark for 30 min, and absorbance was measured at 517 nm using a plate reader. Antioxidant activity was expressed as radical scavenging percentage relative to the control.

##### 2.3.6.2. In vitro antioxidant activity

The *in vitro* antioxidant activity of cloudberry NVs was evaluated in HDF and HaCaT cells using an H_2_O_2_-induced oxidative stress model. HDF and HaCaT cells were seeded in 48-well plates at densities of 3 × 10^3^ and 5 × 10^3^ cells/well, respectively. After overnight attachment, the cells were pretreated with cloudberry NVs at 2, 5, or 10 µg based on protein content for 24 h. The treatment medium was then removed and replaced with culture medium containing 350 µM H_2_O_2_ and SYTOX Green high-affinity nucleic acid stain diluted 1:10,000 to label dead or dying cells. The plates were imaged every 3 h using the IncuCyte live-cell imaging system at 10× magnification under standard scan settings. The samples were analyzed in triplicate.

##### 2.3.6.3. Polyphenol content and related ROS scavenging activity measurement

Total polyphenol content (TPC) of cloudberry NV extracts was determined using the Folin–Ciocalteu method. A gallic acid calibration curve was prepared by dissolving gallic acid (Sigma Aldrich, Merck KGaA, Darmstadt, Germany) in 100% methanol to obtain a stock solution of 10 mg L⁻¹. From this stock, standard solutions of 5, 10, 25, 50, 75, 100, 200 µg mL⁻¹ were prepared. For each standard, 20 µL of solution was mixed with 145 µL of Milli-Q water, 5 µL of Folin–Ciocalteu reagent (Sigma Aldrich, Merck KGaA, Darmstadt, Germany), and 30 µL of sodium carbonate solution. The mixtures were incubated for 45 min in the dark. After incubation, 200 µL of each reaction mixture was transferred to a 96-well plate, and the absorbance was measured at 765 nm using the Multiskan SkyHigh Microplate Spectrophotometer (Thermo Fisher Scientific, USA). The calibration curve equation used to calculate TPC in the samples was y = 0.0017x + 0.0431 where y represents absorbance and x represents gallic acid concentration (µg mL⁻¹). Sample extracts were treated under the same conditions, and their polyphenol concentrations were calculated from the calibration curve and expressed as µg gallic acid equivalents (GAE) per mL of NV solution, with values representing mean ± standard deviation (SD, n = 3).

Antioxidant activity of cloudberry NV extracts was evaluated using the ABTS (2,2′-azino-bis(3-ethylbenzothiazoline-6-sulfonic acid)) radical scavenging assay. Briefly, 1 mL of ABTS radical cation solution was mixed with 100 µL of each polyphenolic extract. After 2.5 min incubation, 200 µL of each reaction mixture was transferred to a 96-well plate, and the absorbance was measured at 734 nm using the Multiskan SkyHigh Microplate Spectrophotometer (Thermo Fisher Scientific, USA). The percentage of ABTS radical scavenging activity was calculated according to the method reported by Ali et al. ^14^, with values representing mean ± standard deviation (SD, n = 3).

### 2.4. Molecular cargo

#### 2.4.1. Proteomic analysis

##### 2.4.1.1. Total protein extraction from cloudberry fruit

Total protein extraction was performed based on the method reported by Stanly et al. ^15^, with minor modifications. Briefly, cloudberry fruit was frozen in liquid nitrogen and homogenized into a fine powder using a blender, with additional liquid nitrogen added between blending steps. Powdered fruit material (2 g) was mixed with 12 mL of extraction buffer containing 2% SDS, 60 mM dithiothreitol (DTT), 20% glycerol, and 40 mM Tris-HCl, pH 8.5, and incubated at 90°C for 8 min. The mixture was then centrifuged at 8,000 × g for 15 min at 4°C using a fixed-angle rotor. The supernatant was collected, mixed with 25 mL of precipitation solution containing 10% w/v trichloroacetic acid and 20 mM DTT in ice-cold acetone, incubated at −20°C for 45 min, and centrifuged at 18,000 × g for 10 min at 4°C. The resulting pellet was washed three times with washing solution containing 20 mM DTT in ice-cold acetone, resuspended in washing solution, incubated at −20°C for 1 h, and centrifuged again at 20,000 × g for 10 min at 4°C. After air-drying for 5 min, the pellet was dissolved in 700 μL of rehydration buffer containing 7 M urea, 2 M thiourea, 30 mM Tris-HCl, pH 8.5, and 43 mM DTT under constant shaking for 1 h at room temperature. Finally, the sample was centrifuged at 12,000 × g for 10 min at room temperature, and the supernatant was collected as the total protein extract.

##### 2.4.1.2. Sample preparation for proteomic analysis

NVs and total protein extracts from cloudberry fruit were subjected to in-solution digestion followed by mass spectrometry (MS) analysis. Samples were lysed in 0.2% RapiGest detergent (Waters Corporation, Milford, MA, USA.) using five freeze–thaw cycles in liquid nitrogen and sonication. Following lysis, proteins were digested with trypsin at a trypsin-to-protein ratio of 1:50. The resulting peptides were purified and enriched using Pierce™ C18 Spin Columns (Thermo Fisher Scientific, USA). Prior to MS analysis, the samples were dissolved in 2% acetonitrile containing 0.1% (v/v) formic acid. MS analysis was performed using a high-resolution hybrid quadrupole time-of-flight mass spectrometer (Waters SELECT SERIES Cyclic IMS, Waters Corp., Wilmslow, UK) equipped with a low-flow electrospray ionization source and coupled to a Waters ACQUITY I-Class UPLC system. Ion mobility separation was used in single pass. Data analysis was performed using ProteinLynx Global SERVER 3.0.3 with closest possible proteome *Rubus argutus* (taxonomy Id: 59490).

##### 2.4.1.3 Proteomics functional analysis

To reduce redundancy associated with the use of the automatically annotated UniProt TrEMBL database, functional characterization of identified proteins was performed using homology-based annotation approaches. Protein sequences were annotated using eggNOG-mapper^16^ for orthology-based functional inference and InterProScan^17^ for conserved domain and Gene Ontology (GO)^18^ assignment. Annotation and downstream GO mapping were implemented through a custom Snakemake^19^-based workflow developed in-house and made publicly available through GitHub^20^. This strategy enabled consolidation of homologous protein annotations and improved functional interpretation of proteomic datasets generated from the non-model plant species. Following annotation, GO assignments obtained from eggNOG-mapper and InterProScan were consolidated using custom Python scripts to generate a non-redundant protein-level functional annotation dataset. To avoid inflation of enrichment statistics caused by repeated homologous annotations, duplicate GO assignments and redundant homolog entries were removed, and a single consolidated GO profile was retained for each protein. Proteins annotated exclusively as hypothetical, predicted, or uncharacterized proteins were excluded from downstream functional interpretation. GO term descriptions and ontology categories were assigned using the Gene Ontology reference database (go-basic.obo).

Gene Ontology overrepresentation analysis (ORA) was performed separately for each sample using Fisher’s exact test^21^ implemented in Python. Because quantitative abundance measurements may not be reliable due to type of database we used for protein identification, enrichment analysis was based on protein presence/absence data. The background universe consisted of all unique proteins detected across both nanovesicle and total protein extract datasets that contained at least one GO annotation, thereby representing the experimentally detectable proteome. P-values were adjusted for multiple testing using the Benjamini–Hochberg false discovery rate^22^ (FDR) correction.

To improve biological interpretability, significantly enriched GO terms were further refined using a custom thematic filtering workflow. Broad and non-informative ontology terms were removed and remaining enriched GO categories were grouped into biologically relevant themes including oxidative stress response, vesicle trafficking, cell protection, cell proliferation, and defense/immunity based on curated keyword classification. Redundant GO descriptions within each thematic category were subsequently collapsed to retain representative biological functions.

Functional annotation confidence was further assessed by integrating Gene Ontology (GO) assignments derived from both eggNOG and InterProScan analyses. Since functional annotations in non-model plant species rely primarily on homology-based inference, GO terms were categorized according to the level of supporting evidence. GO terms identified independently by both eggNOG orthology-based annotation and InterProScan domain-based annotation were considered high-confidence annotations. GO terms identified exclusively through InterProScan were classified as medium confidence due to the presence of conserved functional domains, whereas GO terms assigned only through eggNOG orthology transfer were considered low confidence. Confidence categories were incorporated into downstream overrepresentation analysis and visualization to facilitate interpretation of functionally enriched pathways.

Enrichment results were visualized as bubble plots generated in Python using Matplotlib23, where fold enrichment values were plotted against enriched GO categories and bubble size represented enrichment significance based on −log10(FDR) values.

#### 2.4.2. Metabolomic analysis

##### 2.4.2.1 Extract preparation

Cloudberry NV preparation (250 µL) was mixed with 750 µL of 80% methanol (MeOH). The mixture was vortexed for 1 min and centrifuged at 12,000 rpm for 10 min at 4°C. The resulting supernatant was collected, vacuum-dried, and resuspended in 400 µL of 80% MeOH. Finally, the solution was filtered and stored at −20°C until further analysis. All procedures were performed in triplicate.

##### 2.4.2.2 Untargeted metabolomic analysis

All chemicals and reagents used were either analytical, LC–MS or high-performance liquid chromatography (HPLC) grade reagents. LC-HRMS experiments were performed using a Dionex Ultimate 3000 system equipped with a degasser, quaternary pump, column oven, and autosampler, coupled to a high-resolution Orbitrap mass spectrometer (Q-Exactive, Thermo Fisher Scientific, Waltham, MA, USA). The LC-MS method was carried out on a Kinetex 2.6 μ POLAR C18 100 Å (100 × 3 mm) column (Phenomenex, Torrance, CA, USA), using 0.1 % V/V of HCOOH in H_2_O (solvent A) and acetonitrile (solvent B) as mobile phase. The volume injected was set at 5 µL. The gradient elution was optimized as follows: 5% B for 3 min, increased from 5% to 50% B over 12 min, held for 2 min, then increased from 50% to 95% B over 10 min, held for 3 min, followed by re-equilibration under initial conditions for 5 min. The total run time was 35 min. MS and MS/MS spectra were acquired in both positive and negative ionization modes using data-dependent acquisition (DDA), with fragmentation triggered for the five most intense ions in each full scan. The following settings were used in the full-ion MS mode: resolution power of 70,000 Full Width at Half Maximum (FWHM), scan range 100– 1700 m/z, automatic gain control (AGC) target 1 × 10^6^, injection time set to 100 ms, and scan rate set at 2 scan/s. The ion source parameters were as follows: spray voltage 3.7 kV; capillary temperature 320 °C; S-lens RF level 50; sheath gas pressure 32; auxiliary gas 10; and auxiliary gas heater temperature 350 °C. For the scan event of ddMS2, the parameters were set as follows: mass resolving power = 17,500 FWHM; maximum injection time = 50 ms; AGC target = 1 × 10^5^; scan range = 200–2000 m/z; isolation window 4.0 m/z; and retention time 10 s. The collision energy was set to 25, 30, and 35 eV. The ‘Analysis Base File’ (.abf) format of raw data files were generated using Reifys Abf converter software (https://www.reifycs.com/AbfConverter), a traditional data format for MS-DIAL MS^E^ data processing. MS-DIAL v5.0 software was used for deconvolution, peak picking, alignment, and compound identification ^24^. The detailed parameter settings were as follows: MS1 tolerance, 0.002 Da; MS2 tolerance, 0.002 Da; minimum peak height, 100000 amplitudes; mass slice width, 0.05 Da; smoothing method, linear weighted moving average; smoothing level, 4 scans; minimum peak width, 5 scans; alignment retention time tolerance 0.5 and MS1 tolerance 0.002. Compounds were annotated with accurate precursor masses and MS/MS spectra against all the libraries available on https://systemsomicslab.github.io/compms/msdial/main.html.

### 2.5. Statistical Analysis

For the anti-senescence assays, statistical differences were evaluated using one-way analysis of variance (ANOVA), followed by Šídák’s multiple-comparisons test. For the cellular uptake experiments, quantitative data are presented as the mean ± standard deviation (SD) from three microscopy fields acquired from the same slide per condition. Differences among the 2, 8, and 24 h time points were assessed using Brown–Forsythe and Welch one-way ANOVA, followed by Dunnett’s T3 multiple-comparisons test. A p value < 0.05 was considered statistically significant. Statistical analyses were performed using GraphPad Prism version 10.6.1.

For metabolomics, the raw feature lists, from both positive and negative ionization mode, were subjected to quality control filtering based on the following criteria: well-defined name and formula, MS^2^ fragments, sample-to-blank ratio >5; signal-to-noise (S/N) ratio >10; and missing value fill =50%. Resulting high-quality features (Table S6) were utilized for downstream analysis. Compound classes were identified using ClassyFire database ^25^. Quantitative distribution of major chemical classes was visualized via pie charts generated in R (v.4.1.0) using the *dplyr* (v2.5.1) ^26^ and *ggplot2* (v4.0.2) ^27^ packages. Pathway enrichment analysis was performed by importing the filtered feature list into MetaboAnalyst v6.0 ^28^, utilizing *Arabidopsis thaliana* as the reference plant species for metabolic mapping (Table S7). The experimental workflow (Fig. 7A) was conceptualized and drafted with the assistance of Google Gemini 3.1 Pro (Google, 2026). The final figure was reviewed and edited by the authors.

## 3. Results

### 3.1. Physicochemical characterization of cloudberry NVs

The physicochemical properties of cloudberry NVs were in agreement with those reported previously for this vesicle population ^12^. NTA analysis confirmed a nanosized vesicle preparation with a mean particle diameter of around 200-250 nm, and TEM imaging revealed characteristic round vesicular structures. Protein quantification by Qubit assay together with particle counting by NTA demonstrated a measurable recovery of NV-associated material from cloudberry fruit (from 1 kg of fruit 13 ± 1.4 mg of protein, and 6.2*10^12^ ± 1.2*10^11^ particles). Although the protein yield was different than previously reported result, ^12^ particle yield remains in the same range, suggesting cloudberry fruit as a suitable source for isolation of PDNVs.

### 3.2. Cytotoxicity and cell proliferation

The effects of cloudberry NVs on cell viability, proliferation, and cytotoxicity were assessed in HDF and HaCaT cells by MTT assay and IncuCyte live-cell imaging. In both cell lines, the response was concentration-dependent, with lower doses being better tolerated than the highest dose. In the MTT assay, treatment with 2 and 5 µg NVs resulted in a moderate reduction in viability compared with the control, whereas 10 µg caused a more pronounced decrease in both HDF and HaCaT cells (Fig. 1A). Similarly, IncuCyte confluence analysis showed a dose-dependent reduction in cell proliferation, with the strongest inhibitory effect observed at 10 µg (Fig. 1B). SYTOX Green-based cytotoxicity monitoring, presented as normalized green object count, further supported these trends.

**Figure 1.**
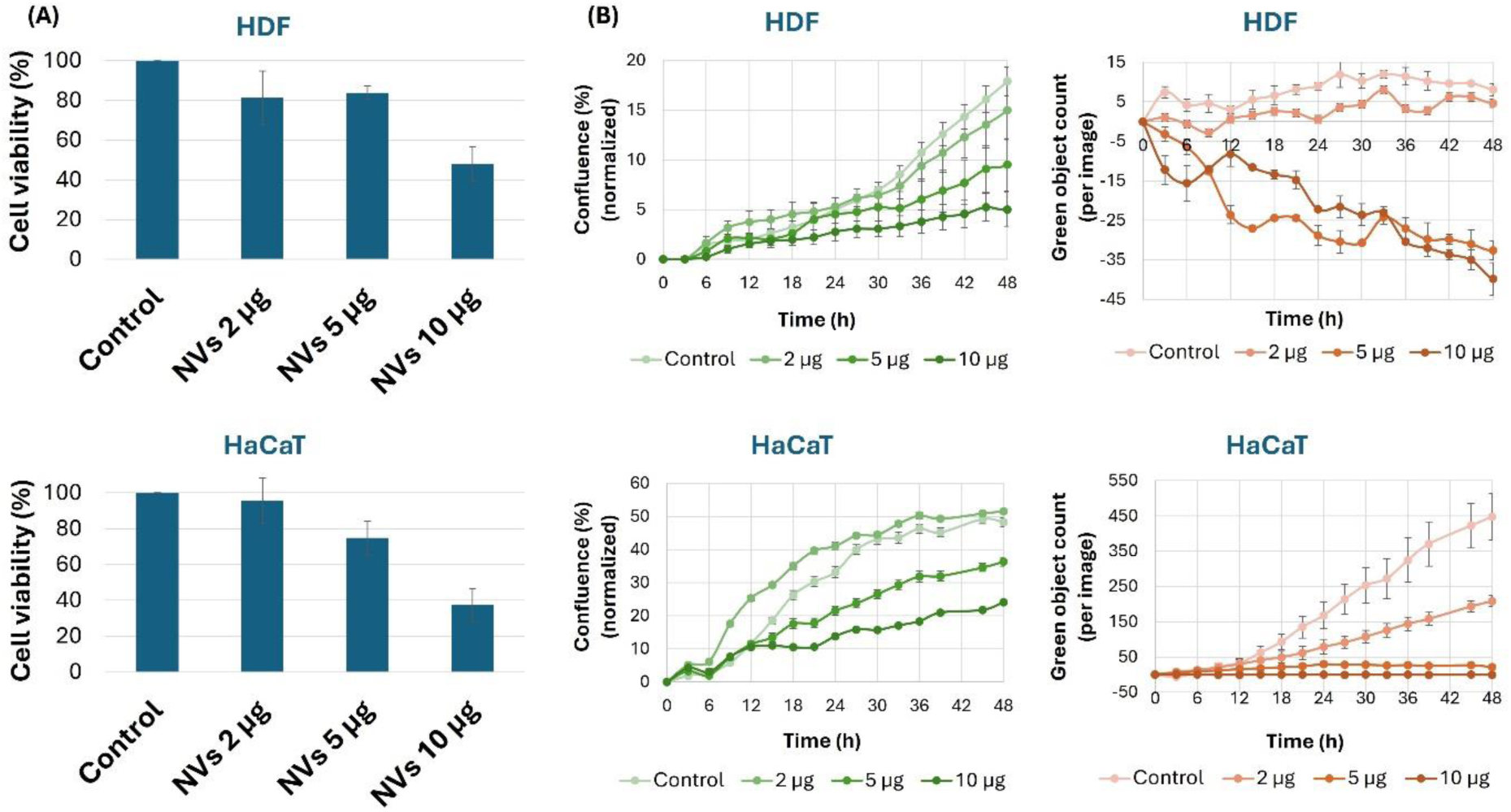
Dose-dependent effects of cloudberry NVs on cell proliferation and cytotoxicity in HDFs and HaCaT cells, assessed by (A) MTT assay and (B) Incucyte live-cell imaging.

In HaCaT cells, the control group showed the highest increase in normalized green object count over time, whereas NV-treated groups remained markedly lower. In HDF cells, NV treatment also altered the normalized green object count profile in a concentration-dependent manner, with the clearest deviation from the control observed at 10 µg. Our results indicate that cloudberry NVs modulated cell viability and growth in a dose-dependent manner, with 2–5 µg being relatively better tolerated than 10 µg.

### 3.3. Cellular uptake of cloudberry NVs by the cells

Confocal microscopy revealed time-dependent uptake of DiOC18-labelled cloudberry NVs by both HDF and HaCaT cells (Fig. 2A). In both cell types, the green fluorescence signal was relatively weak and sparse after 2 h, increased after 8 h, and was most pronounced after 24 h of incubation. Quantification of the background-corrected green fluorescence intensity per cell confirmed a significant time-dependent increase in NV-associated fluorescence in both HDF and HaCaT cells, with the highest values detected at 24 h (Fig. 2B).

**Figure 2.**
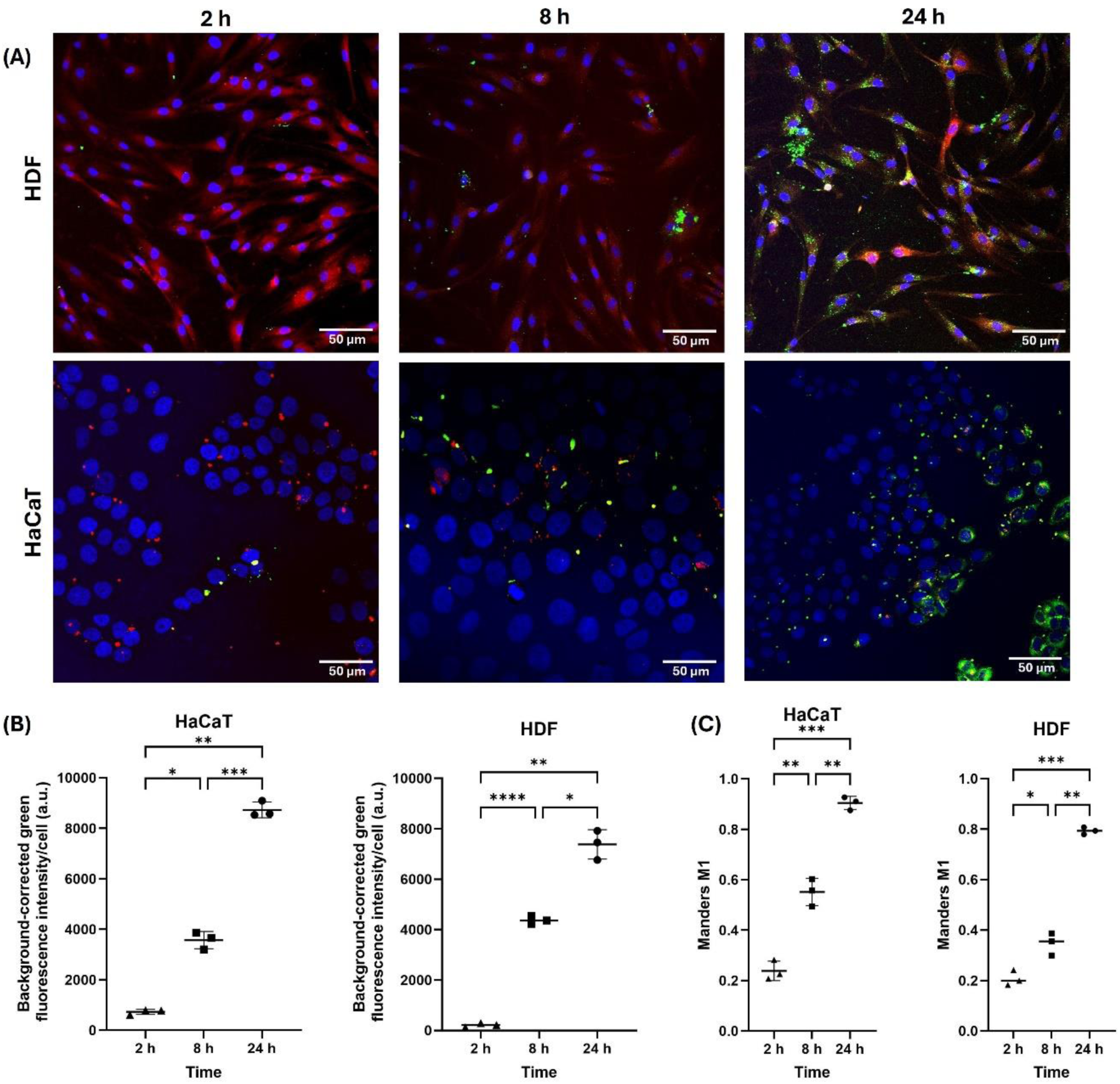
Time-dependent uptake and lysosomal colocalization of cloudberry-derived nanovesicles in HDF and HaCaT cells. (A) Representative confocal microscopy images of human dermal fibroblasts (HDFs) and HaCaT keratinocytes after 2, 8, and 24 h of incubation with cloudberry-derived nanovesicles (NVs). DiOC18-labelled NVs are shown in green, lysosomes stained with LysoTracker Red are shown in red, and Hoechst-stained nuclei are shown in blue. (B) Quantification of the background-corrected green fluorescence intensity per cell as a measure of NV uptake. (C) Quantification of NV–lysosome colocalization using Manders’ M1 coefficient. Data are presented as mean ± SD. Statistical significance is indicated as p < 0.05, *p < 0.01, **p < 0.001, and ***p < 0.0001.

The NV-associated green fluorescence increasingly overlapped with the LysoTracker-stained lysosomal signal over time. Accordingly, Manders’ M1 coefficient increased significantly from 2 to 24 h in both cell types, reaching approximately 0.9 in HaCaT cells and 0.8 in HDFs at 24 h (Fig. 2C). These findings indicate progressive cellular uptake of cloudberry NVs and their subsequent accumulation within, or trafficking towards, the endolysosomal compartment. Both dermal fibroblasts and keratinocytes efficiently internalized cloudberry NVs, with the greatest uptake and lysosomal colocalization observed after 24 h.

### 3.4. Anti-senescent activity of cloudberry NVs

The anti-senescent potential of cloudberry NVs was evaluated in HDF cells using senescence-associated β-galactosidase staining after H_2_O_2_-induced stress. As expected, treatment with H_2_O_2_ markedly increased the number of SA-β-gal-positive cells compared with the untreated control, indicating successful induction of a senescence-like phenotype (Fig. 3A). In contrast, cells treated with cloudberry NVs alone at 2, 5, or 10 µg showed only minimal SA-β-gal staining, similar to the control. Pretreatment with cloudberry NVs before H_2_O_2_ exposure reduced the extent of positive staining compared with the H_2_O_2_-treated group, suggesting a protective effect against oxidative stress-induced cellular senescence. Quantification of SA-β-gal-positive cells confirmed these observations, showing a significant increase after H₂O₂ treatment and a significant reduction in the 5 µg + H₂O₂ and 10 µg + H₂O₂ groups compared with H₂O₂ alone, while the 2 µg + H₂O₂ group did not show a significant reduction (Fig. 3B). Among the tested concentrations, the 5 µg condition appeared to show the clearest reduction in senescence-associated staining, whereas protection was less evident at 2 and 10 µg. Overall, cloudberry NVs exert anti-senescent activity in HDF cells under oxidative stress conditions.

**Figure 3.**
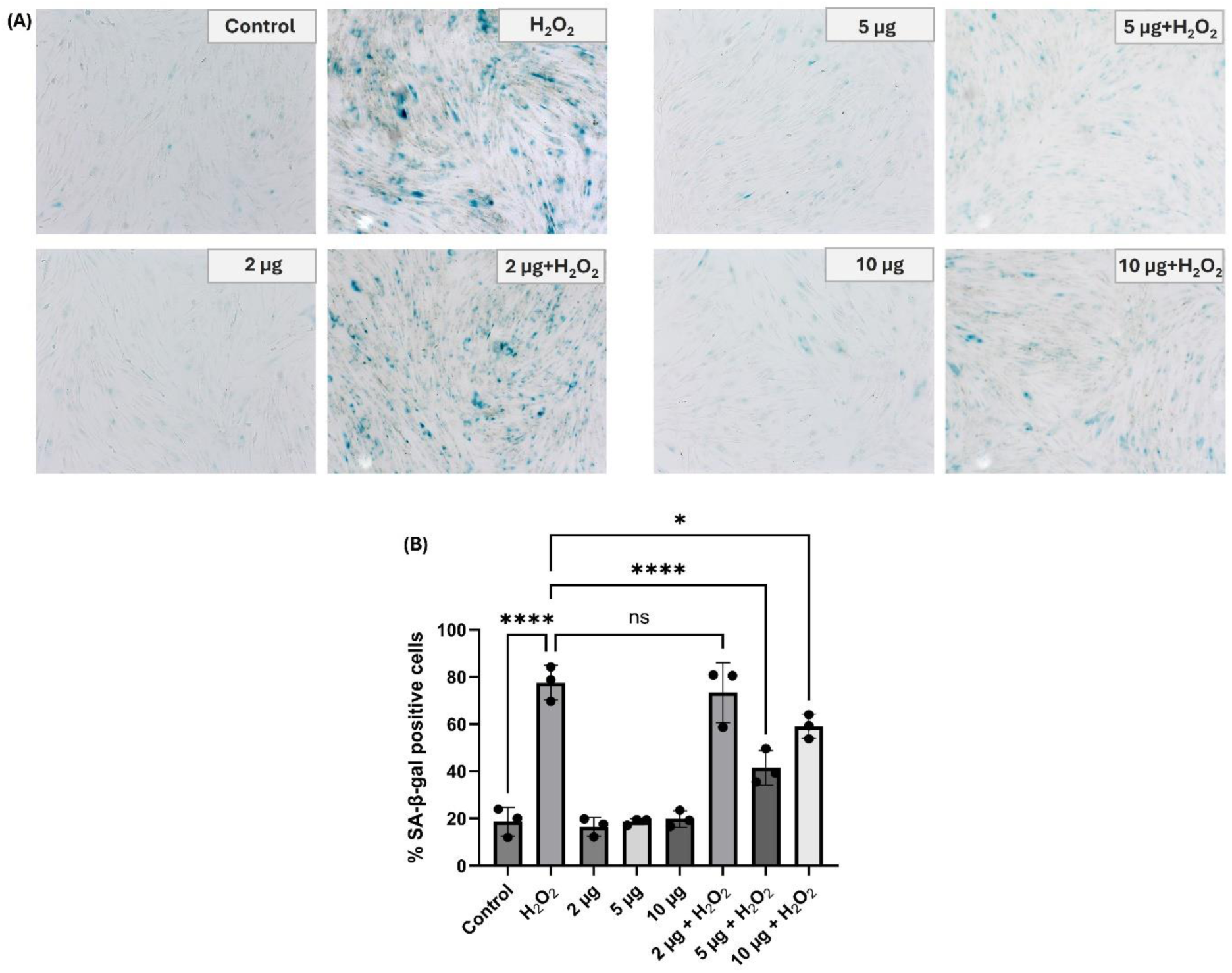
(A) Cloudberry NVs prevent H2O2-induced cellular senescence in HDF cells. Positive SA-β-gal staining indicates senescent cells. Scale bars, 100 µm; (B) Percentage of SA-β-Gal-positive cells (n = 3). *p < 0.05, ****p < 0.0001.

### 3.5. Wound-healing activity of cloudberry NVs

The effect of cloudberry NVs on wound closure was evaluated in HDF and HaCaT cells using a scratch wound-healing assay monitored with the IncuCyte system for 48 h (Fig. 4). In both cell types, cloudberry NVs influenced wound closure in a dose-dependent manner. In HDF cells, treatment with 2 µg NVs accelerated wound closure compared with the untreated control during the early and intermediate time points, whereas 5 µg showed a weaker effect and 10 µg markedly inhibited wound closure throughout the observation period (Fig. 4C,E). Wound-width measurement consistently revealed that the control and 2 µg group’s wound width gradually decreased, almost completely closing after 48 hours, however the 5 µg group’s reduction was slower and the 10 µg group’s wound width remained high throughout the experiment (Fig. 4E). By 48 h, both the control and 2 µg groups reached a similarly high degree of closure. In HaCaT cells, the control and 2 µg groups showed the fastest wound closure overall, while 5 µg delayed closure and 10 µg showed minimal wound healing activity during the experiment (Fig. 4D,F). This pattern was further supported by wound width measurements, where the control and 2 µg groups showed a consistent decrease in wound width, to almost zero values, while the 5 µg groups resulted in delayed wound closure, and wound width of 10 µg treatment group remained noticeably larger (Fig. 4F). Representative images at 24 and 48 h were consistent with the quantitative analysis, showing substantial narrowing of the wound area in the control and 2 µg groups, a slower response at 5 µg, and limited closure at 10 µg in both HDF and HaCaT cells (Fig. 4A,B). These results indicate that cloudberry NVs modulate wound-healing-related cell migration in a dose-dependent manner, with 2 µg showing the most favorable effect in HDF cells, while in HaCaT cells its effect was broadly comparable to the untreated control.

**Figure 4.**
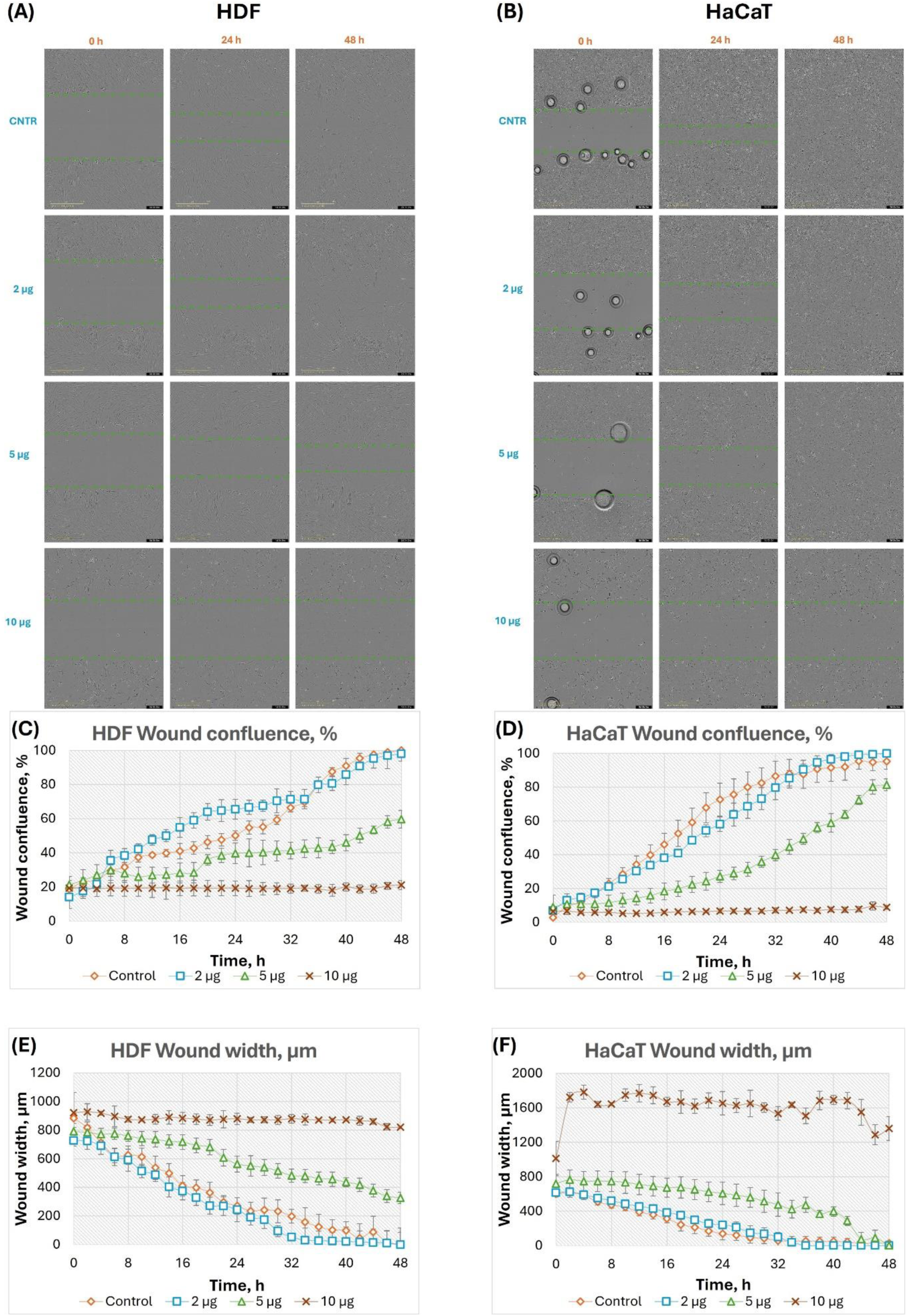
Effects of cloudberry NVs on wound closure in HDF and HaCaT cells. Wound closure was monitored for 48 h after treatment with cloudberry NVs at 2, 5, or 10 µg based on protein content, or without treatment (control). Representative images of (A) HDF and (B) HaCaT cells at 0, 24, and 48 h are shown. Quantitative analysis is presented as wound confluence over time for (C) HDF and (D) HaCaT cells, and as wound width over time for (E) HDF and (F) HaCaT cells, recorded using the IncuCyte S3 live-cell analysis system.

### 3.6. Antioxidant activity

The antioxidant potential of cloudberry NVs was first assessed using the DPPH radical scavenging assay. As shown in Fig. 5A, cloudberry NVs exhibited measurable radical scavenging activity at all tested doses, with values remaining in a similar range across 2, 5, and 10 µg, indicating that the vesicle preparation possesses cell-free antioxidant capacity. This effect appeared slightly higher at 5 µg, although the overall differences between doses were modest.

**Figure 5.**
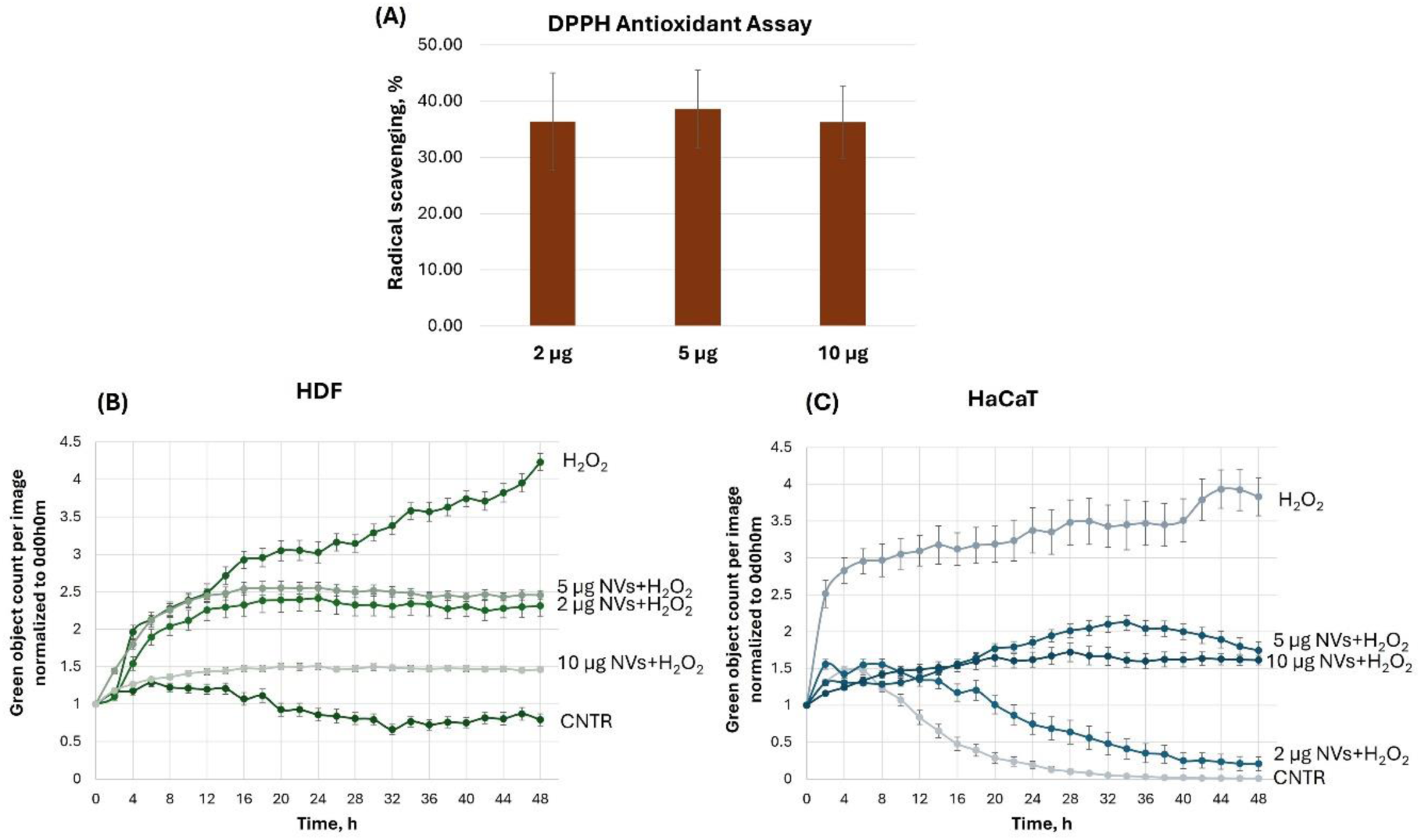
Antioxidant activity of cloudberry NVs. (A) Cell-free antioxidant activity measured by the DPPH radical scavenging assay at 2, 5, and 10 µg of NVs. (B) In vitro antioxidant activity in HDF cells and (C) HaCaT cells under H2O2-induced oxidative stress.

The antioxidant activity of cloudberry NVs was further evaluated in HDF and HaCaT cells under H_2_O_2_-induced oxidative stress conditions using SYTOX Green-based live-cell imaging. In both cell lines, treatment with H_2_O_2_ alone markedly increased the green object count over time compared with the untreated control, confirming induction of oxidative stress-associated cell damage (Fig. 5B,C). Pretreatment with cloudberry NVs attenuated this response in a dose-dependent manner, although the pattern differed between the two cell types. In HDF cells, the strongest protective effect was observed at 10 µg, which maintained the green object count closest to the control level throughout the experiment, whereas 2 and 5 µg showed only partial protection. In HaCaT cells, the most pronounced protective effect was observed at 2 µg, which reduced the signal almost to the control level by the later time points, while 5 and 10 µg produced a more moderate reduction compared with the H_2_O_2_-treated group. These findings indicate that cloudberry NVs exert antioxidant and cytoprotective effects against H_2_O_2_-induced oxidative damage, although the optimal dose appears to be cell type-dependent.

### 3.7. Proteomic analysis

To further investigate proteins that may contribute to the antioxidant and wound-healing properties of cloudberry-derived nanovesicles and total protein extract (PE), both samples were subjected to proteomic analysis. Protein identification was performed against the UniProtKB/TrEMBL *Rubus argutus* database. A total of 1041 proteins were identified in NVs, whereas 619 proteins were identified in the PE (Supplementary Tables S1–S3). Among these, 415 proteins were shared between the two samples, while 626 proteins were uniquely detected in NVs (Fig. 6A).

**Figure 6.**
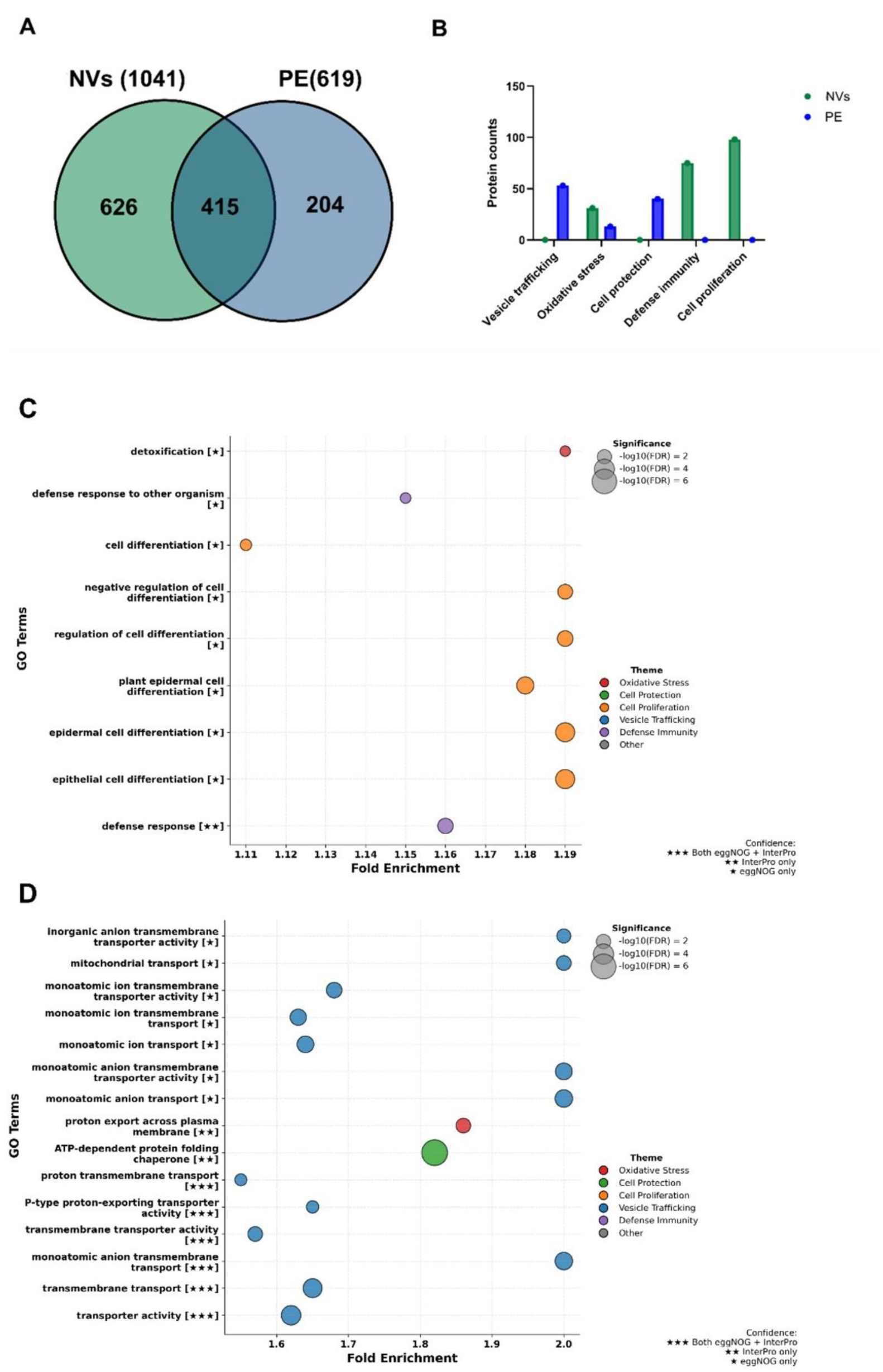
Proteomic and functional characterization of cloudberry-derived nanovesicles and protein extract (PE). (A) Venn diagram showing the overlap and unique distribution of proteins identified in NVs and PE. (B) Bar plot illustrating the major biological themes assigned to enriched Gene Ontology (GO) categories and the number of proteins associated with each theme. (C) Bubble plot representing enriched GO categories identified in NV proteome. (D) Bubble plot representing enriched GO categories identified in the PE proteome. Bubble size corresponds to enrichment significance [−log10(FDR)], while the x-axis indicates fold enrichment values. GO annotation confidence is indicated by star symbols, where ★★★ represents GO terms supported by both eggNOG and InterProScan, ★★ represents InterProScan-supported annotations and ★ represents eggNOG-derived annotations.

To improve functional characterization and minimize redundancy associated with automatically annotated TrEMBL entries, homolog-based functional annotation was performed using eggNOG-mapper and InterProScan. Functional annotations generated from both platforms were integrated using a custom bioinformatics workflow to obtain non-redundant protein-level Gene Ontology (GO) assignments. As the analysis was conducted in a non-model plant species, the resulting GO annotations were interpreted as homology-supported functional predictions.

Gene Ontology overrepresentation analysis (ORA), followed by thematic filtering of GO categories associated with oxidative stress response, cell survival, and wound-healing-related processes, revealed clear functional differences between the two samples (Fig. 6B–D). NVs were enriched in proteins associated with oxidative stress response, defence and immunity, and cell proliferation-related processes, whereas the PE showed greater enrichment for proteins involved in vesicle trafficking and cellular protection pathways (Fig. 6B–D).

Proteins associated with oxidative stress-related GO annotations were predominantly linked to glutathione-dependent and thioredoxin-mediated redox metabolism pathways (Table 1). Several independent antioxidant-associated systems were represented within the NV proteome, including glutathione transferases, glutathione peroxidases, glutathione reductase, thioredoxin-related proteins, catalase, superoxide dismutase, and peroxiredoxin-associated proteins. Collectively, these proteins are involved in cellular redox homeostasis, detoxification of reactive oxygen species (ROS), and adaptation to oxidative stress.

**Table 1.** Proteins associated with oxidative stress response and antioxidant-related GO terms in cloudberry-derived nanovesicles.

| Protein Name | Protein IDs |
| --- | --- |
| Catalase | A0AAW1WVR1 |
| Glutaredoxin-dependent peroxiredoxin | A0AAW1X9C2 |
| Glutathione peroxidase | A0AAW1WT33, A0AAW1WU93 |
| Glutathione reductase (GRase) | A0AAW1X306 |
| Glutathione S-transferase | A0AAW1XL78, A0AAW1XLH6, A0AAW1XM57, A0AAW1XMR8, A0AAW1XNT0, A0AAW1XNU5, A0AAW1XNZ0, A0AAW1XPT2 |
| Glutathione transferase | A0AAW1WHK6, A0AAW1WHX0, A0AAW1XMH6, A0AAW1XNS5, A0AAW1YQ70 |
| Monodehydroascorbate reductase (NADH) | A0AAW1WCT9 |
| Protein-disulfide reductase | A0AAW1XG40, A0AAW1XGJ3 |
| Superoxide dismutase [Cu-Zn] | A0AAW1W754 |
| Thioredoxin domain-containing protein | A0AAW1W2P7, A0AAW1W7I8, A0AAW1XP98, A0AAW1YD51, A0AAW1YHA4 |
| Thioredoxin reductase | A0AAW1YI87 |
| Thioredoxin-dependent peroxiredoxin | A0AAW1WD36 |
| Uncharacterized protein | A0AAW1XAK1, A0AAW1XC40 |

In contrast, proteins associated with cell proliferation-related GO annotations were primarily represented by homeobox domain-containing and START domain-containing proteins (Table 2). These proteins are generally associated with developmental regulation, cellular organization, membrane-associated signalling, and intracellular transport processes. START domain-containing proteins have additionally been linked to lipid transfer and membrane dynamics, which are important for vesicle-mediated cellular communication.

**Table 2.** Proteins contributing to cell proliferation-associated GO enrichment in cloudberry-derived nanovesicles.

| Protein Name | Protein IDs |
| --- | --- |
| AAA+ ATPase domain-containing protein | A0AAW1Y348 |
| Acetyl-CoA C-acetyltransferase | A0AAW1WLZ6 |
| Actin | A0AAW1WT39 |
| CRAL-TRIO domain-containing protein | A0AAW1X098 |
| DNA topoisomerase 6 subunit B | A0AAW1XXI0 |
| Dynamin GTPase | A0AAW1XYG1 |
| Glycerophosphodiester phosphodiesterase | A0AAW1XTA3 |
| Guanosine nucleotide diphosphate dissociation inhibitor | A0AAW1YEW6 |
| Homeobox domain-containing protein | A0AAW1VK81, A0AAW1VR71, A0AAW1VRQ6, A0AAW1VUR6, A0AAW1VXC9, A0AAW1W4I5, A0AAW1WDL3, A0AAW1WDY6, A0AAW1WFK5, A0AAW1WVS3, A0AAW1WWM9, A0AAW1WWS8, A0AAW1WXG6, A0AAW1WZ44, A0AAW1X0E9, A0AAW1X789, A0AAW1X7U5, A0AAW1X9Y1, A0AAW1XTL9, A0AAW1XUN7, A0AAW1Y1P9, A0AAW1Y352, A0AAW1Y376, A0AAW1Y461, A0AAW1YP35, A0AAW1YQA7, A0AAW1YS37, A0AAW1YS83 |
| Non-specific lipid-transfer protein | A0AAW1W9T4 |
| Patellin-3 | A0AAW1X280 |
| Phosphopyruvate hydratase | A0AAW1WN97 |
| Protein-disulfide reductase | A0AAW1XG40, A0AAW1XGJ3 |
| Ras-related protein Rab7 | A0AAW1Y442 |
| S-adenosylmethionine synthase | A0AAW1WAF0 |
| Serine hydroxymethyltransferase | A0AAW1VMZ4, A0AAW1XB32 |
| START domain-containing protein | A0AAW1VF65, A0AAW1VHI4, A0AAW1VKP4, A0AAW1VLF5, A0AAW1VRJ5, A0AAW1VZ41, A0AAW1W052, A0AAW1WVT1, A0AAW1WW90, A0AAW1WXY5, A0AAW1WZF6, A0AAW1WZR1, A0AAW1X6B4, A0AAW1X9F5, A0AAW1XU38, A0AAW1Y459, A0AAW1Y4F1, A0AAW1Y4Z1, A0AAW1Y565, A0AAW1Y5I3, A0AAW1YSD2 |
| Uncharacterized protein | A0AAW1VKN6, A0AAW1VLD4, A0AAW1VPX2, A0AAW1VQH9, A0AAW1VWJ3, A0AAW1VXY9, A0AAW1W2W0, A0AAW1W9D8, A0AAW1WEB7, A0AAW1WER0, A0AAW1WF59, A0AAW1WFX4, A0AAW1WI34, A0AAW1WLM9, A0AAW1WT77, A0AAW1WVE3, A0AAW1WWB9, A0AAW1WXB6, A0AAW1WYM9, A0AAW1WYS2, A0AAW1WZP6, A0AAW1X1G0, A0AAW1X5T1, A0AAW1X6V3, A0AAW1XB96, A0AAW1XKQ9, A0AAW1XRY3, A0AAW1XSJ4, A0AAW1Y7C3, A0AAW1YRB6 |
| Uridine kinase | A0AAW1WNX9 |
| Vignain | A0AAW1WRN6 |

As oxidative stress regulation and controlled redox signalling are important components of tissue repair and cellular regeneration, the enrichment of proteins associated with antioxidant defence, cellular protection, and growth-associated regulatory pathways suggests that protein cargo of NVs may have contributed to the oxidative stress management and wound-healing-related cellular processes observed in the experimental data.

### 3.8. Metabolomic analysis

To characterize the chemical composition of cloudberry-derived nanovesicles, metabolites were extracted and subjected to comprehensive profiling (Fig. 7A). Given that cloudberries are known reservoirs of bioactive polyphenols, we first quantified the total polyphenol content (TPC). The NVs exhibited a TPC of 33.74 μg GAE/mL (Fig. S1), suggesting a significant encapsulation of polar and semi-polar compounds. Consistent with this polyphenol enrichment, the extracts demonstrated potent biological activity, achieving 65.60% inhibition of reactive oxygen species (ROS) formation (Fig. S1).

**Figure 7.**
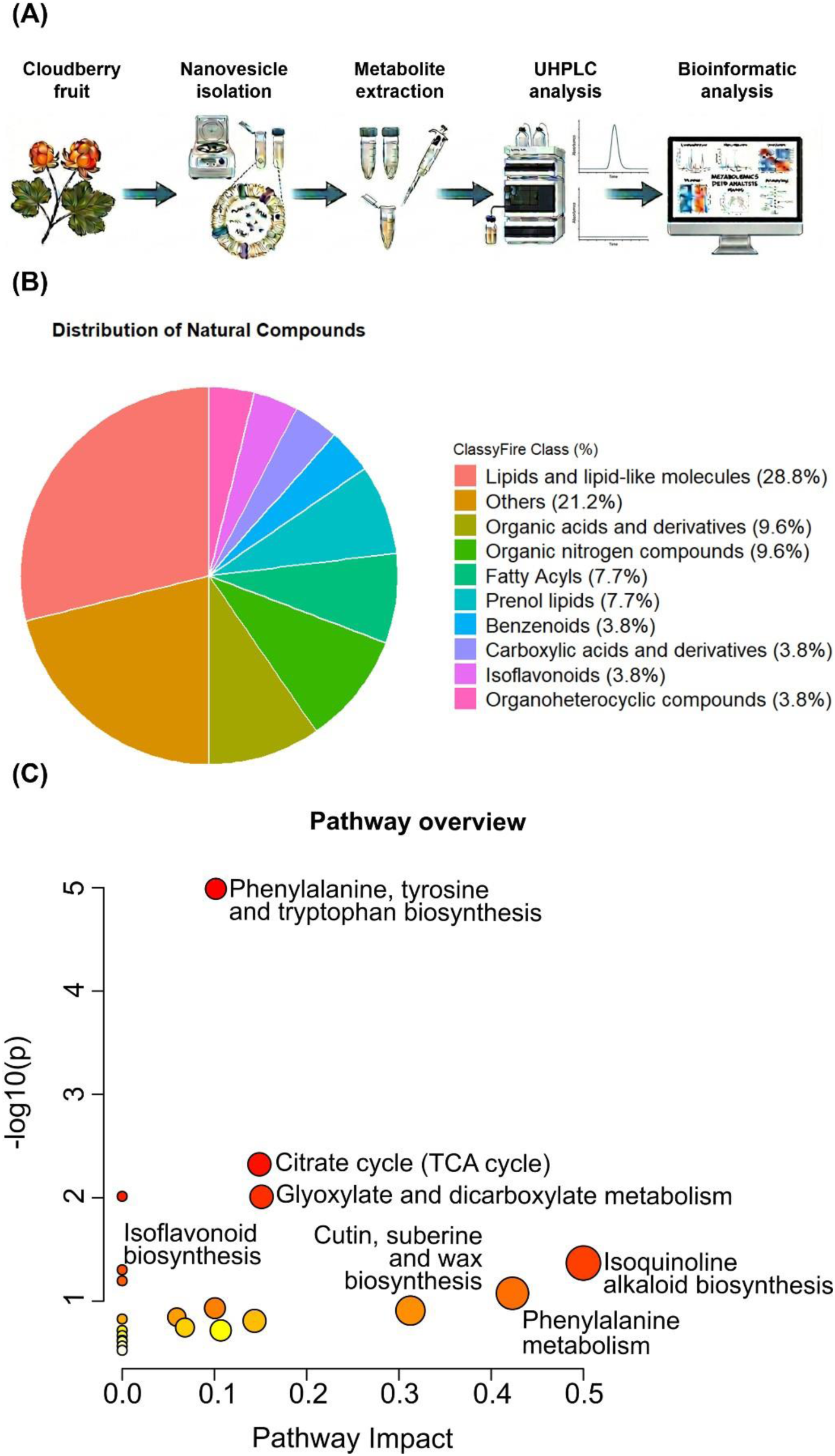
Metabolomic characterization of cloudberry-derived NVs. (A) Methodological pipeline illustrating NV isolation, metabolite extraction, UHPLC-MS/MS analysis, and bioinformatic interpretation. (B) Pie chart representing the distribution of the most abundant compound classes in NVs, expressed as a percentage of the total identified chemical classes. (C) Scatter plot of pathway enrichment analysis in NVs. Each circle represents an individual metabolic pathway; the x-axis indicates pathway impact, and the y-axis represents the significance level (-log10(p)). Circle size is proportional to pathway impact, while the color gradient (yellow to red) reflects increasing statistical significance.

Subsequent untargeted metabolomic analysis performed in both negative and positive ionization modes (Fig. S2A and B) revealed the accumulation of 53 compounds within the NVs (Table S6). Classification of the detected features showed that lipids and lipid-like molecules (28.8%) and organic acids and organic nitrogen compounds (9.6%) constituted the primary chemical framework of the vesicles (Fig. 7B). Notably, prenol lipids (7.7%) and isoflavonoids (3.8%) frequently associated with the health-promoting properties of fruits— were well represented (Fig. 7B).

To provide biological context to these findings, a pathway enrichment analysis was conducted. The metabolic profile of the NVs was significantly centered on the biosynthesis and metabolism of phenylalanine, tyrosine, and tryptophan (Fig. 7C).

Furthermore, distinct enrichment was observed across primary and secondary metabolic pathways, specifically the citrate (TCA) cycle, glyoxylate and dicarboxylate metabolism, and isoflavonoid biosynthesis (Fig. 7C).

In accordance with these results, the 17 most representative metabolites identified within the NVs—assigned at confidence levels 2 or 3 based on MS/MS spectral library matching—are presented in Table 3. These included 6 polyphenols, 6 terpenoids, and 5 lipids/sphingolipids. Among the polyphenols, ellagic acid was the most prominent, exhibiting a peak area of 1.4×10^9^. These findings correlate with the identification of amino acids (phenylalanine, L-tryptophan, and leucine) and primary metabolites (citric, malic, and shikimic acids) (Table S7), further supporting the activation of phenylalanine and tryptophan biosynthetic pathways and the TCA cycle. These data suggest that NVs transport a diverse molecular cargo encompassing both primary and secondary metabolic intermediates.

**Table 3.**
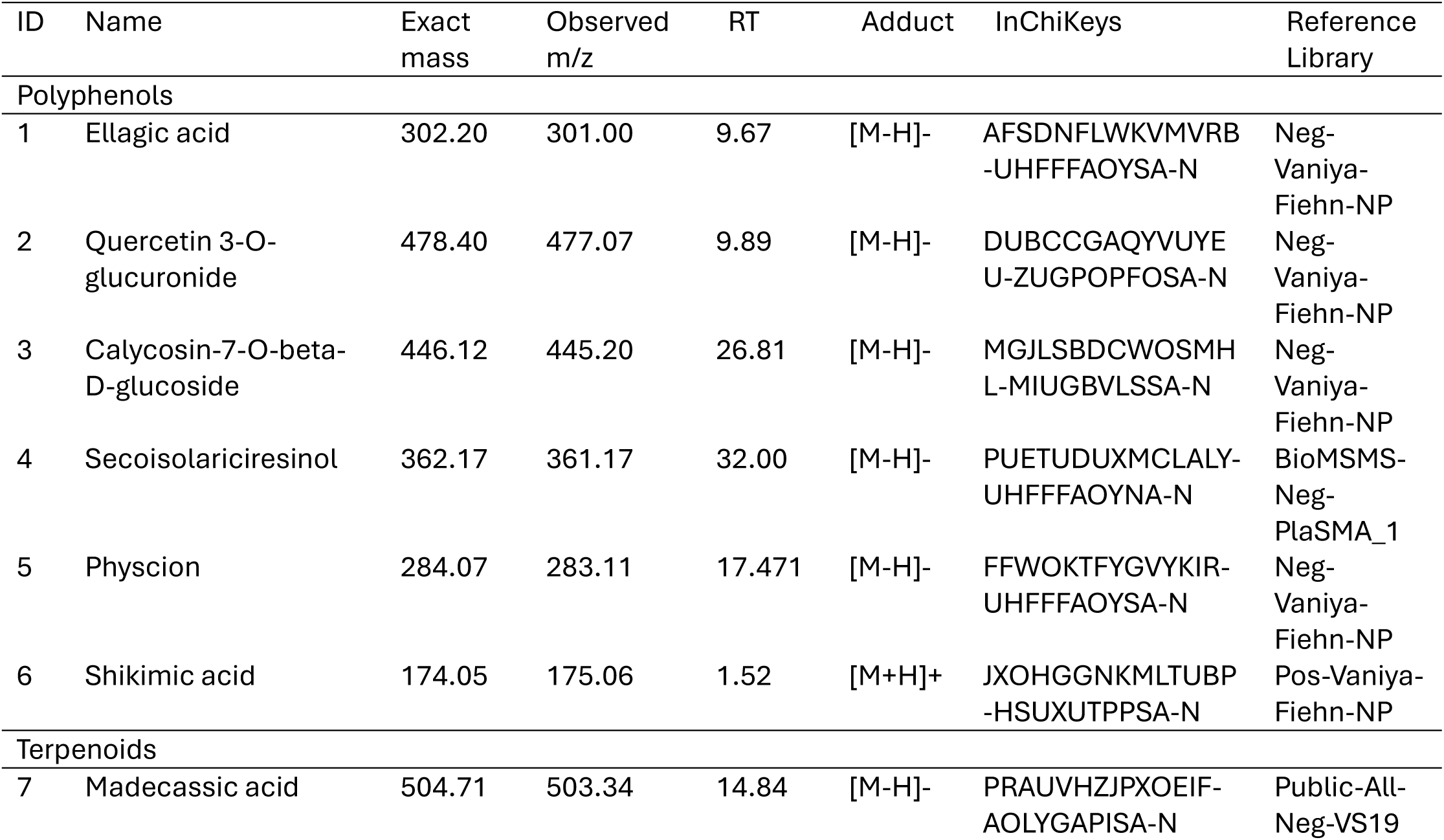
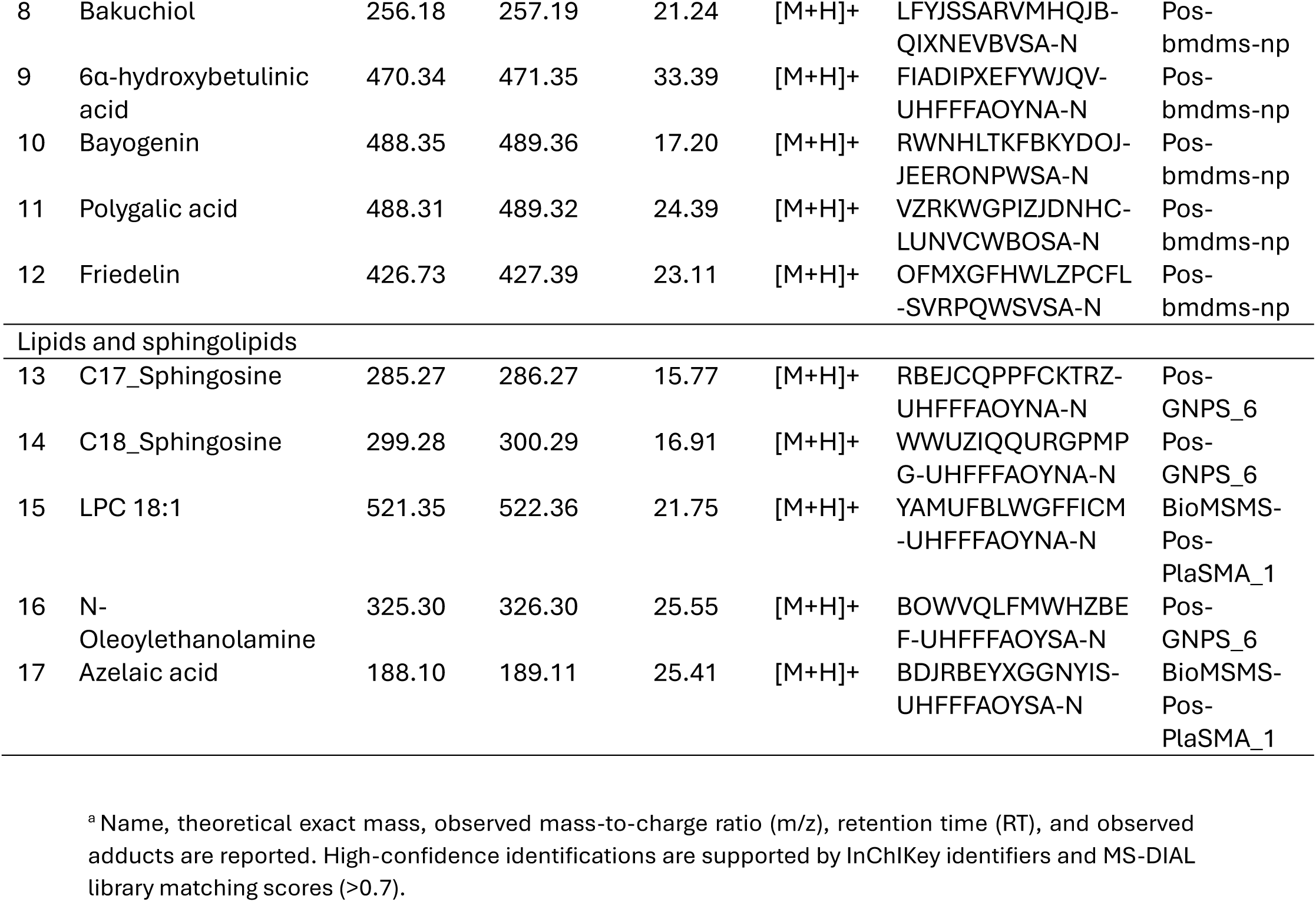
Putative metabolites annotated by matching their fragmentation spectra to public databases.

Furthermore, several triterpenoids were notably accumulated, including bayogenin (C_30_H_46_O_5_, 3.25×10^7^), friedelin (C_30_H_50_O, 1.63×10^7^), 6α-hydroxybetulinic acid (C_30_H_46_O_4_, 5.63×10^7^), and polygalic acid (C_29_H_44_O_6_, 1.40×10^7^) (Table 3). The presence of lipids and sphingolipids was also confirmed, specifically the accumulation of C17_ and C18_ sphingosines and LPC 18:1. Collectively, the presence of these diverse metabolic classes, ranging from primary amino acids and organic acids to bioactive specialized metabolites, suggests that cloudberry-derived NVs serve as a complex transport vehicle for both biosynthetic precursors and final antioxidant products.

## 4. Discussion

Cloudberry-derived nanovesicles have promising physicochemical properties for anti-aging applications. Their nanoscale size and negative surface charge may support interactions with skin-relevant cells, however, further studies are required to evaluate their penetration and delivery behavior in skin models. In addition, their natural lipid bilayer can protect bioactive antioxidant metabolites and proteins from degradation, thereby preserving their stability and functional activity. Moreover, the findings of this study indicate that cloudberry-derived NVs are not only physically stable vesicle-like particles, but also biologically active systems capable of interacting with skin-relevant cells. In vitro experiments showed that cloudberry-derived NVs are biologically and functionally active, and interact with skin-relevant cells in a concentration-dependent manner. In both HDF and HaCaT cells, lower doses were generally better tolerated than 10 µg, whereas the highest dose reduced viability/proliferation and impaired functional responses. This non-linear behavior is important because recent plant-vesicle studies show that beneficial effects on skin cells are strongly dependent on vesicle source, dose, and recipient cell type. For example, tomato-derived nanovesicles promoted migration of keratinocytes and fibroblasts, ginseng-derived nanoparticles enhanced proliferation and migration of HaCaT, BJ fibroblast-like cells and HUVECs, and mixed plant-derived EV preparations showed antioxidant and reparative activity in human skin fibroblasts ^7,29,30^. Therefore, the present data suggest that cloudberry NVs may have a useful biological window, with low-to-intermediate concentrations being more suitable for skin-related applications than higher concentrations.

The time-dependent uptake of labelled cloudberry NVs by HDF and HaCaT cells supports their potential role as carriers of bioactive plant-derived cargo. The stronger fluorescent signal after 24 h, together with partial colocalization with lysosomal staining in HDFs, suggests internalization followed by intracellular trafficking, at least for a fraction of the labelled vesicle-associated material. Comparable uptake by skin-relevant cells has been reported for tomato-and ginseng-derived nanovesicles, where internalization was linked to enhanced migration, proliferation, or repair-related responses ^29,30^. The ability of plant-derived vesicles to transport molecular cargo, including proteins, lipids, metabolites, and RNAs, is considered one of the main reasons for their biological effects in recipient cells ^3,4,11,31,32^. At the same time, lipophilic membrane dyes can generate dye aggregates or label non-vesicular components; therefore, the dye-only controls used here are essential, and future studies should include complementary strategies such as quantitative flow cytometry, cargo tracking, or labelled vesicle proteins/RNAs to confirm vesicle-specific uptake.

A major functional finding was the antioxidant and cytoprotective effect of cloudberry NVs under H_2_O_2_-induced stress. The DPPH assay showed intrinsic radical-scavenging capacity, while IncuCyte-based monitoring demonstrated reduced oxidative stress-associated cell damage in both HDF and HaCaT cells. These results agree with recent reports showing that plant-derived vesicles can mitigate oxidative injury in skin models. Di Raimo et al. reported antioxidant and reparative effects of a plant-derived EV mix in human skin fibroblasts ^7^, while ginseng root-derived exosome-like nanoparticles protected skin cells from UV-induced oxidative stress by limiting ROS generation and suppressing AP-1-related signaling ^33^. Ginger-derived EVs also reduced ROS and apoptosis in UVA-induced skin injury models ^34^, and aloe-derived nanoparticles were recently shown to attenuate photoaging-associated oxidative stress and senescence through activation of the Nrf2/ARE pathway ^35^. In this context, the reduction of H_2_O_2_-induced SA-β-gal staining in HDFs suggests that cloudberry NVs may counteract stress-induced senescence-like changes, although the optimal protective dose differed between assays and cell types.

The proteomic profile of cloudberry NVs provides a possible molecular explanation for these antioxidant and cytoprotective effects. The NV fraction contained a broad protein cargo, with a larger number of identified proteins than the total protein extract and a substantial set of proteins uniquely detected in NVs. This is consistent with previous proteomic studies showing that plant EVs and plant-derived vesicle preparations are enriched in proteins related to stress responses, defense, redox regulation, and vesicle trafficking ^36–38^. For example, Rutter and Innes showed that Arabidopsis apoplastic EVs carry stress-response proteins, including proteins associated with oxidative stress and defense responses ^36^. Similarly, recent reviews of plant EV proteomes have reported the presence of HSP70, annexins, phospholipases, glutathione S-transferases, and other proteins involved in ROS signaling and stress adaptation ^38^.

Functional annotation and GO-based analysis in the present study indicated enrichment of proteins related to oxidative stress response, defense/immunity, and cell proliferation-related processes in cloudberry NVs. In particular, the presence of antioxidant-associated proteins such as glutathione transferases, glutathione peroxidases, glutathione reductase, thioredoxin-related proteins, catalase, superoxide dismutase, peroxiredoxin-associated proteins, and monodehydroascorbate reductase supports the idea that cloudberry NVs may carry protein systems involved in redox homeostasis and ROS detoxification. These protein groups are well known components of plant antioxidant defense, and their presence in EV-like preparations supports the possibility that cloudberry NVs contribute to oxidative stress buffering in recipient cells ^36,38^. Therefore, these proteins may act together with vesicle-associated polyphenols and triterpenoids to explain the observed protection against H_2_O_2_-induced cell damage and the reduction of senescence-associated β-galactosidase staining.

The wound-healing assay further showed that cloudberry NVs modulate cell migration/closure in a dose-dependent manner. In HDFs, 2 µg accelerated wound closure during the early and intermediate phases, whereas 10 µg clearly delayed closure; in HaCaT cells, the lowest dose behaved similarly to the control, while higher doses impaired closure. This pattern differs from some pro-migratory plant vesicle studies, such as tomato-derived nanovesicles, ginseng-derived nanoparticles, and ginger-derived EVs, which promoted migration and/or proliferation in skin-relevant cells ^29,30,34^. However, it is consistent with the cytocompatibility data in the present work and indicates that excessive cloudberry NV exposure may suppress proliferation or alter cell metabolism sufficiently to slow scratch closure. Because scratch assays reflect both migration and proliferation, future experiments should separate these processes using proliferation-controlled migration assays and include the analysis of migration and extracellular matrix remodeling-related markers, together with inflammatory mediators. In addition to these functional observations, the investigation of cloudberry-derived NVs revealed a diverse metabolome with potent antioxidant and health-promoting potential, mirroring findings in nanovesicles from plants such as *Salvia miltiorrhiza* ^39^ and *Cynara cardunculus* L. var. altilis ^40^, as well as various fruits ^7^. However, our untargeted metabolomic analysis further explored possible links between the chemical cargo of cloudberry NVs and their observed biological activity. Our identification of ellagic acid is in strong agreement with existing literature. Guo et al. ^41^ recently demonstrated that cloudberries are particularly enriched in tannins with high ellagic acid content compared to related berries. This highlights the capacity of cloudberry NVs to naturally sequester ellagic acid, a compound with well-documented antioxidant, anti-inflammatory, and wound-healing properties ^42,43^. Notably, while previous studies showed that EVs engineered to encapsulate ellagic acid could promote diabetic wound healing and collagen formation without reducing skin cell viability ^44^, our results demonstrate that cloudberry NVs are naturally enriched with this metabolite, underscoring their inherent therapeutic value. The proteomic data also support the wound-healing-related observations, as cloudberry NVs contained proteins associated with cell proliferation, membrane trafficking, cytoskeletal organization, lipid transfer, and vesicle dynamics, including homeobox domain-containing proteins, START domain-containing proteins, actin, dynamin GTPase, Rab7, non-specific lipid-transfer protein, and patellin-3. Although these findings do not prove direct activation of wound-healing pathways, they suggest that cloudberry NV cargo may contribute to cell–vesicle interactions and the dose-dependent effects observed in the scratch assay, consistent with previous reports identifying trafficking-and signaling-related proteins in plant EV proteomes ^37^.

Furthermore, the accumulation of triterpenoids within these NVs is of pivotal importance. Among the triterpenic acids identified, friedelin was previously detected at high levels in cloudberry by Falev et al. ^45^. Triterpenic acids are renowned for their antioxidant properties ^46^. For instance, betulinic acid has been shown to reduce senescence in human dermal fibroblasts ^47^. This aligns with our finding of significant 6α-hydroxybetulinic acid accumulation. Triterpenic acids carried by EVs can demonstrate an important antioxidant potential as shown for EVs from *Salvia* hairy roots, which were able to neutralize neurotoxic molecules and protect SH-SY5Y neuroblastoma cells ^48^.

Finally, our untargeted analysis revealed a distinct enrichment in lipids and sphingolipids, including C17_ and C18_sphingosine and lysophosphatidylcholine (LPC) 18:1. These molecules are essential for both structural integrity and inter-kingdom signaling. While C18_sphingosine serves as a fundamental membrane component, lysolipids like LPC 18:1 facilitate membrane curvature during biogenesis and have been implicated in modulating host immune responses following ingestion ^31,32^.

Taken together, the functional, metabolomic, and proteomic data suggest that the biological activity of cloudberry NVs is likely mediated by combined cargo, including antioxidant polyphenols, triterpenoids, bioactive lipids, and proteins related to redox regulation, defense responses, vesicle trafficking, and membrane organization. This integrated cargo may explain the antioxidant, anti-senescent, and wound-healing-related effects observed in skin-relevant cells; however, because proteomic annotation was based on homology in a non-model plant species and plant vesicle preparations may contain co-isolated non-vesicular components, these findings should be considered functional indications rather than definitive mechanistic proof, requiring further validation ^1,2,49^.

Overall, our data indicate that cloudberry-derived NVs are biologically active plant vesicles with potential use in skin-related antioxidant, anti-senescent, and wound-healing applications. However, their effects appear to depend on the dose and cell type, emphasising the importance of further investigations to better understand their modes of action, confirm vesicle-specific cargo delivery, and determine the optimal dose range for potential cosmetic or biological applications.

## 5. Conclusion

In this study, nanovesicles containing diverse molecular cargo were isolated from cloudberry fruit and characterized. The results demonstrated that these NVs are internalized by skin cells and modulate several functional responses in vitro, including cell viability, cellular uptake, oxidative stress response, senescence-associated β-galactosidase staining, and wound closure. Lower concentrations were tolerated better, and favoured more biological effects, while the higher concentrations reduced cell proliferation and migration, showing a clear dose-dependent effect. Proteomic analysis revealed proteins associated with redox regulation, defense responses, vesicle trafficking, and growth-related processes, supporting the functional effects observed in skin-relevant cells. Metabolomic analysis further identified polyphenols, triterpenoids, and lipids, including compounds with antioxidant, anti-inflammatory and skin repair related properties. Altogether, these results indicate that cloudberry-derived NVs are promising natural nanosystems with potential for skin-related applications.

## CRediT authorship contribution statement

**Ramila Mammadova:** Investigation, Formal analysis, Methodology, Data curation, Writing – original draft. **Feby Wijaya Pratiwi:** Investigation, Methodology, Formal analysis, Writing – review & editing. **Carmen Laezza**: Writing – review & editing, Formal analysis, Software. **Keerthanaa Balasubramanian Shanthi**: Writing – review & editing, Formal analysis, Software. **Dávid Papp:** Methodology, Formal analysis. **Carmina Sirignano**: Methodology, Formal analysis. **Dilki Madubhashani**: Investigation, Data curation. **Maria Manuela Rigano:** Methodology, Supervision. **Gitta Schlosser**: Methodology, Supervision. **Seppo Vainio**: Supervision, Conceptualization, Funding acquisition, Project administration.

## Data Availability Statement

All data generated during this study are included in this published article and its supplementary information file.

## Acknowledgements

This work was supported by the NutriEV project funded by the European Union under grant agreement no. 101161353. Views and opinions expressed are those of the author(s) only and do not necessarily reflect those of the European Union or the European Innovation Council and SMEs Executive Agency (EISMEA). Neither the European Union nor EISMEA can be held responsible for them. The authors also acknowledge EU/Interreg Aurora (WaVes project grant no. 20370365). Ramila Mammadova gratefully acknowledges the support from the Jenny and Antti Wihuri Foundation for funding her postdoctoral research through the PoDoCo programme and Lumene Oy for the research collaboration. The project was supported by the Lendület (Momentum) Program of the Hungarian Academy of Sciences and by the SNN 148580 project implemented with the support provided by the Ministry of Innovation and Technology of Hungary from the National Research, Development and Innovation Fund, financed under the SNN_24 funding scheme, and by the Slovenian Research and Innovation Agency (ARIS).

The authors acknowledge the contribution of the Biocenter Oulu Light Microscopy Core Facility, a part of Biocenter Finland’s Biological Imaging Platform, and thank Veli-Pekka Ronkainen for his assistance with microscopy image acquisition. The authors also gratefully acknowledge Johanna Kekolahti-Liias and Paula Haipus for their valuable technical assistance.

