## Supplementary Figures for "Cloudberry-derived nanovesicles: *in vitro* functional effects in skin cell models and characterization of molecular cargo"

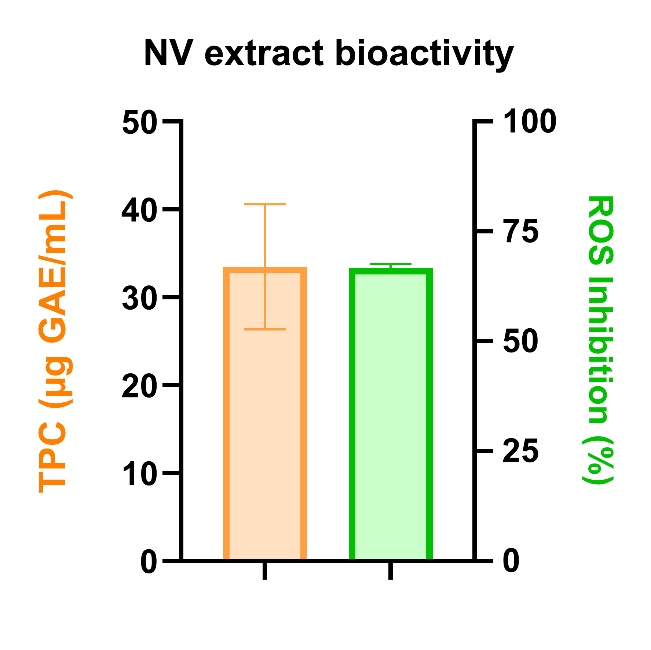

**Figure S1**. Bar graph showing the total polyphenol content (TPC) in ug GAE/mL and the % of ROS inhibition exhibited by the same extract.

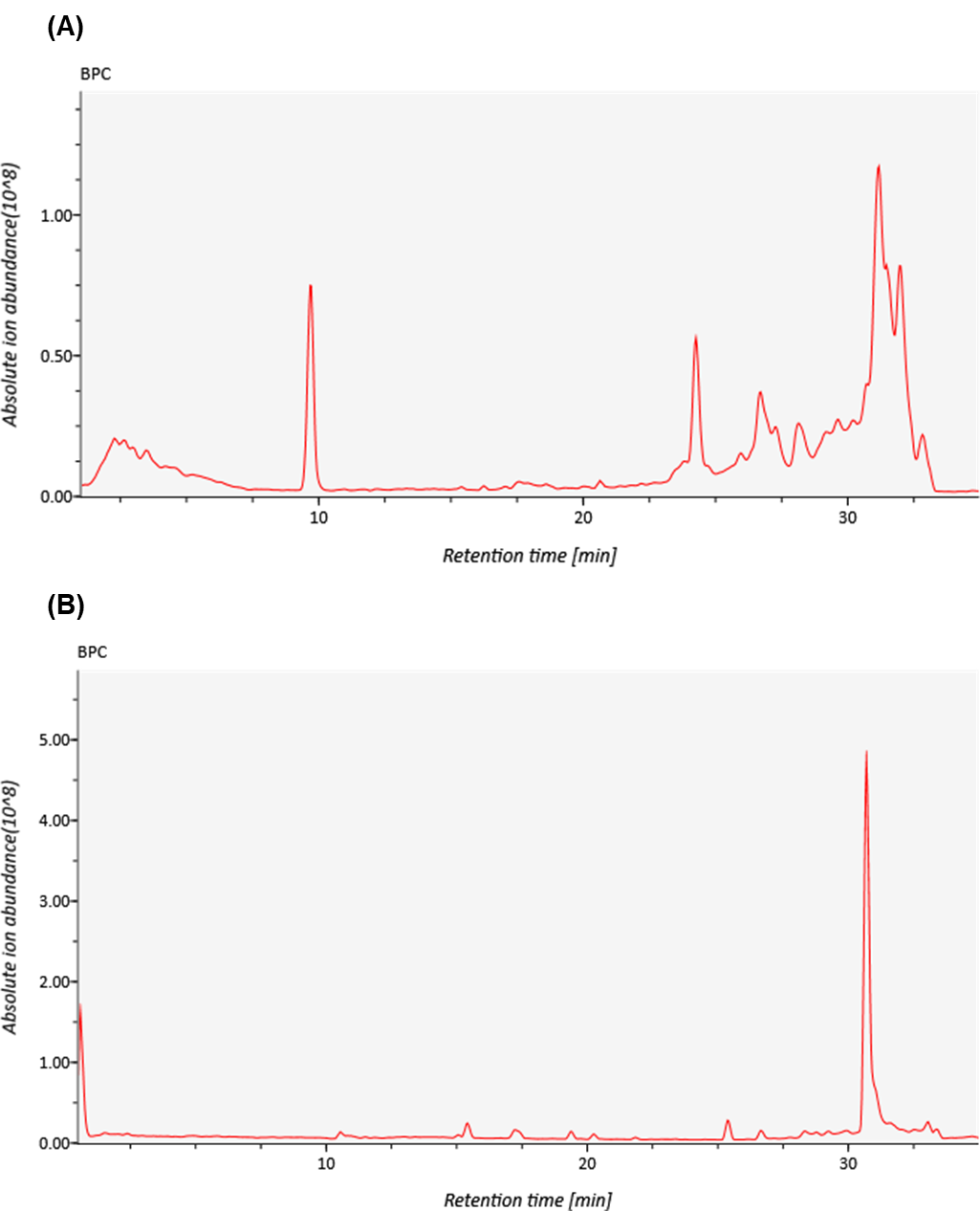

**Figure S2**. Representative Base Peak Chromatograms (BPC) of cloudberry nanovesicle extracts. The metabolic profiles were obtained using UHPLC-MS/MS in both **(A)** negative and **(B)** positive electrospray ionization modes.
